# The human visual system preserves the hierarchy of 2-dimensional pattern regularity

**DOI:** 10.1101/2021.02.05.429884

**Authors:** Peter J. Kohler, Alasdair D. F. Clarke

## Abstract

Symmetries are present at many scales in images of natural scenes. A large body of literature has demonstrated contributions of symmetry to numerous domains of visual perception. The four fundamental symmetries, reflection, rotation, translation and glide reflection, can be combined in exactly 17 distinct ways. These *wallpaper groups* represent the complete set of symmetries in 2D images and have recently found use in the vision science community as an ideal stimulus set for studying the perception of symmetries in textures. The goal of the current study is to provide a more comprehensive description of responses to symmetry in the human visual system, by collecting both brain imaging (Steady-State Visual Evoked Potentials measured using high-density EEG) and behavioral (symmetry detection thresholds) data using the entire set of wallpaper groups. This allows us to probe the hierarchy of complexity among wallpaper groups, in which simpler groups are subgroups of more complex ones. We find that this hierarchy is preserved almost perfectly in both behavior and brain activity: A multi-level Bayesian GLM indicates that for most of the 63 subgroup relationships, subgroups produce lower amplitude responses in visual cortex (posterior probability: > 0.95 for 56 of 63) and require longer presentation durations to be reliably detected (posterior probability: > 0.95 for 49 of 63). This systematic pattern is seen only in visual cortex and only in components of the brain response known to be symmetric-specific. Our results show that representations of symmetries in the human brain are precise and rich in detail, and that this precision is reflected in behavior. These findings expand our understanding of symmetry perception, and open up new avenues for research on how fine-grained representations of regular textures contribute to natural vision.

---

Symmetries are abundant in natural and man-made environments, due to a complex interplay of physical forces that govern pattern formation in nature. Symmetrical patterns have been created and appreciated by human cultures throughout history and since the gestalt movement of the early 20th century, symmetry has been recognized as important for visual perception. Symmetry contributes to the perception of shapes (Palmer, 1985; Li et al., 2013), scenes (Apthorp and Bell, 2015) and surface properties (Cohen and Zaidi, 2013), as well as the social process of mate selection (Møller, 1992). Most of this work has focused on mirror symmetry or *refection*, with much less attention being paid to the other fundamental symmetries: *rotation, translation* and *glide refection*. In the two spatial dimensions relevant for images, these four symmetries can be combined in 17 distinct ways, *the wallpaper groups* (Fedorov, 1891; Polya, 1924; Liu et al., 2010). Previous work on a subset of four of the wallpaper groups used functional MRI to demonstrate that rotation symmetries in wallpapers elicit parametric responses in several areas in occipital cortex, beginning with visual area V3 (Kohler et al., 2016). This effect was also robust with electroencephalography (EEG), whether measured using Steady-State Visual Evoked Potentials (SSVEPs)(Kohler et al., 2016) or event-related paradigms (Kohler et al., 2018). Here we extend this work by collecting SSVEPs and psychophysical data from human participants viewing the full set of wallpaper groups. We measure responses in visual cortex to 16 out of the 17 wallpaper groups, with the 17th serving as a control stimulus. Our goal is to provide a more complete picture of how wallpaper groups are represented in the human visual system.

A wallpaper group is a topologically discrete group of isometries of the Euclidean plane, i.e. transformations that preserve distance (Liu et al., 2010). Wallpaper groups differ in the number and kind of these transformations. In mathematical group theory, when the elements of one group is completely contained in another, the inner group is called a subgroup of the outer group (Liu et al., 2010). The full list of subgroup relationships is listed in Section 1.4.2 of the Supplementary Material. Subgroup relationships between wallpaper groups can be distinguished by their indices. The index of a subgroup relationship is the number of cosets, i.e. the number of times the subgroup is found in the supergroup (Liu et al., 2010). As an example, let us consider groups *P2* and *P6*. If we ignore the translations in two directions that both groups share, group *P6* consists of the set of rotations {0°, 60°, 120°, 180°, 240°, 300°}, in which *P2* {0°, 180°} is contained. *P2* is thus a subgroup of *P6*, and *P6* can be generated by combining *P2* with rotations {0°, 120°, 240°}. Because *P2* is repeated three times in *P6, P2* is a subgroup of *P6* with index 3 (Liu et al., 2010). The 17 wallpaper groups thus obey a hierarchy of complexity where simpler groups are subgroups of more complex ones (Coxeter and Moser, 1972).

The two datasets we present here make it possible to assess the extent to which both behavior and brain responses follow the hierarchy of complexity expressed by the subgroup relationships. Based on previous brain imaging work showing that patterns with more axes of symmetry produce greater activity in visual cortex (Sasaki et al., 2005; Kohler et al., 2018, 2016; Keefe et al., 2018), we hypothesized that more complex groups would produce larger SSVEPs. For the psychophysical data, we hypothesized that more complex groups would lead to shorter symmetry detection thresholds, based on previous data showing that under a fixed presentation time, discriminability increases with the number of symmetry axes in the pattern (Wagemans et al., 1991). Our results confirm both hypotheses, and show that activity in human visual cortex is remarkably consistent with the hierarchical relationships between the wallpaper groups, with SSVEP amplitudes and psychophysical thresholds following these relationships at a level that is far beyond chance. The human visual system thus appears to encode all of the fundamental symmetries using a representational structure that closely approximates the subgroup relationships from group theory.

## Results

The stimuli used in our two experiments were generated from random-noise textures, which made it possible to generate multiple exemplars from each of the wallpaper groups, as described in detail elsewhere (Kohler et al., 2016). We generated control stimuli matched to each exemplar in the main stimulus set, by scrambling the phase but maintaining the power spectrum. All wallpaper groups are inherently periodic because of their repeating lattice structure. Phase scrambling maintains this periodicity, so the phase-scrambled control images all belong to group *P1* regardless of group membership of the original exemplar. *P1* contains no symmetries other than translation, while all other groups contain translation in combination with one or more of the other three fundamental symmetries (reflection, rotation, glide reflection) (Liu et al., 2010). In our SSVEP experiment, this stimulus set allowed us to isolate brain activity specific to the symmetry structure in the exemplar images from activity associated with modulation of low-level features, by alternating exemplar images and control exemplars. In this design, responses to structural features beyond the shared power spectrum, including any symmetries other than translation, are isolated in the odd harmonics of the image update frequency (Kohler et al., 2016; Norcia et al., 2015, 2002). Thus, the combined magnitude of the odd harmonic response components can be used as a measure of the overall strength of the visual cortex response.

The psychophysical experiment took a distinct but related approach. In each trial an exemplar image was shown with its matched control, one image after the other, and the order varied pseudo-randomly such that in half the trials the original exemplar was shown first, and in the other half the control image was shown first. After each trial, participants were instructed to indicate whether the first or second image contained more structure. The duration of both images was controlled by a staircase procedure so that a threshold duration for symmetry detection could be computed for each wallpaper group.

Examples of the wallpaper groups and a summary of our brain imaging and psychophysical measurements are shown in Figure 1. For our primary SSVEP analysis, we only considered EEG data from a pre-determined region-of-interest (ROI) consisting of six electrodes over occipital cortex (see Supplementary Figure 1.1). SSVEP data from this ROI was filtered so that only the odd harmonics that capture the symmetry response contribute to the waveforms. While waveform amplitude is quite variable among the 16 groups, all groups have a sustained negative-going response that begins at about the same time for all groups, 180 ms after the transition from the *P1* control exemplar to the original exemplar. To reduce the amplitude of the symmetry-specific response to a single number that could be used in further analyses and compared to the psychophysical data, we computed the root-mean-square (RMS) over the odd-harmonic-filtered waveforms. The data in Figure 1 are shown in descending order according to RMS. The psychophysical results, shown in box plots in Figure 1, were also quite variable between groups, and there seems to be a general pattern where wallpaper groups near the top of the figure, that have lower SSVEP amplitudes, also have longer psychophysical threshold durations.

**Figure 1:**
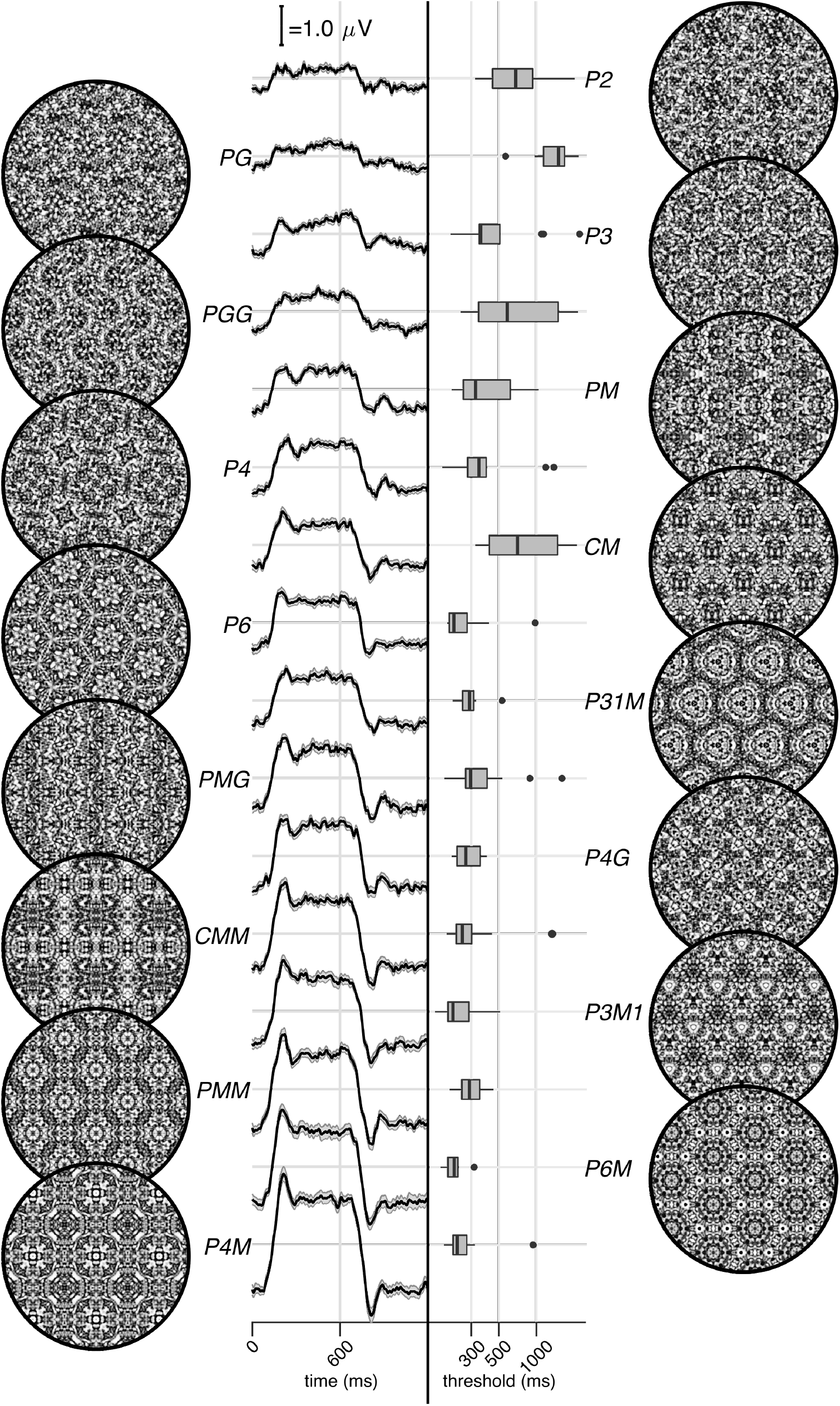
Examples of each of the 16 wallpaper groups are shown in the left- and right-most column of the figures, next to the corresponding SSVEP (center-left) and psychological (center-right) data from each group. The SSVEP data are odd-harmonic-fìltered cycle-average waveforms. In each cycle, a *P1* exemplar was shown for the first 600 ms, followed by the original exemplar for the last 600 ms. Errorbars are standard error of the mean. Psychophysical data are presented as boxplots reflecting the distribution of display duration thresholds. The 16 groups are ordered by the strength of the SSVEP response, to highlight the range of response amplitudes.

We now wanted to test our two hypotheses about how SSVEP amplitudes and threshold durations would follow subgroup relationships, and thereby quantify the degree to which our two measurements were consistent with the group theoretical hierarchy of complexity. We tested each hypothesis using the same approach. We first fitted a Bayesian model with wallpaper group as a factor and participant as a random effect. We fit the model separately for SSVEP RMS and psychophysical data and then computed posterior distributions for the difference between supergroup and subgroup. These difference distributions allowed us to compute the conditional probability that the supergroup would produce (a) larger RMS and (b) a shorter threshold durations, when compared to the subgroup. The posterior distributions are shown in Figure 2 for the SSVEP data, and in Figure 3 for the psychophysical data, which distributions color-coded according to conditional probability. For both data sets our hypothesis is confirmed: For the overwhelming majority of the 63 subgroup relationships, supergroups are more likely to produce larger symmetry-specific SSVEPs and shorter symmetry detection threshold durations, and in most cases the conditional probability of this happening is extremely high.

**Figure 2:**
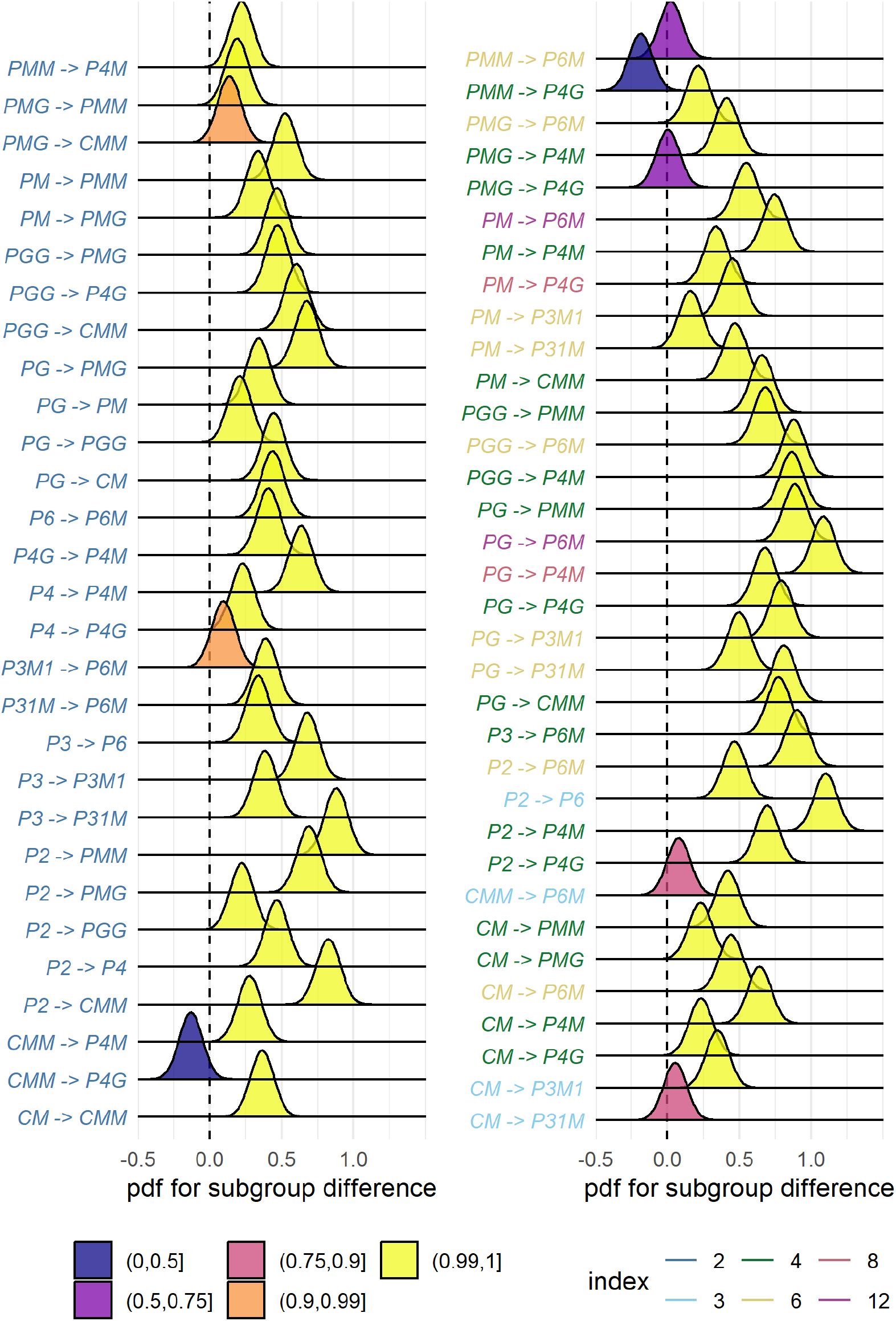
Posterior distributions for the difference in mean SSVEP RMS amplitude. Colour coding of the text indicates the index of the subgroup, while the colour of the filled distribution relates to the conditional probability that the difference in means is greater than zero. We can see that 55/63 subgroup relationships have *p*(Δ|*data*) > 0.99.

**Figure 3:**
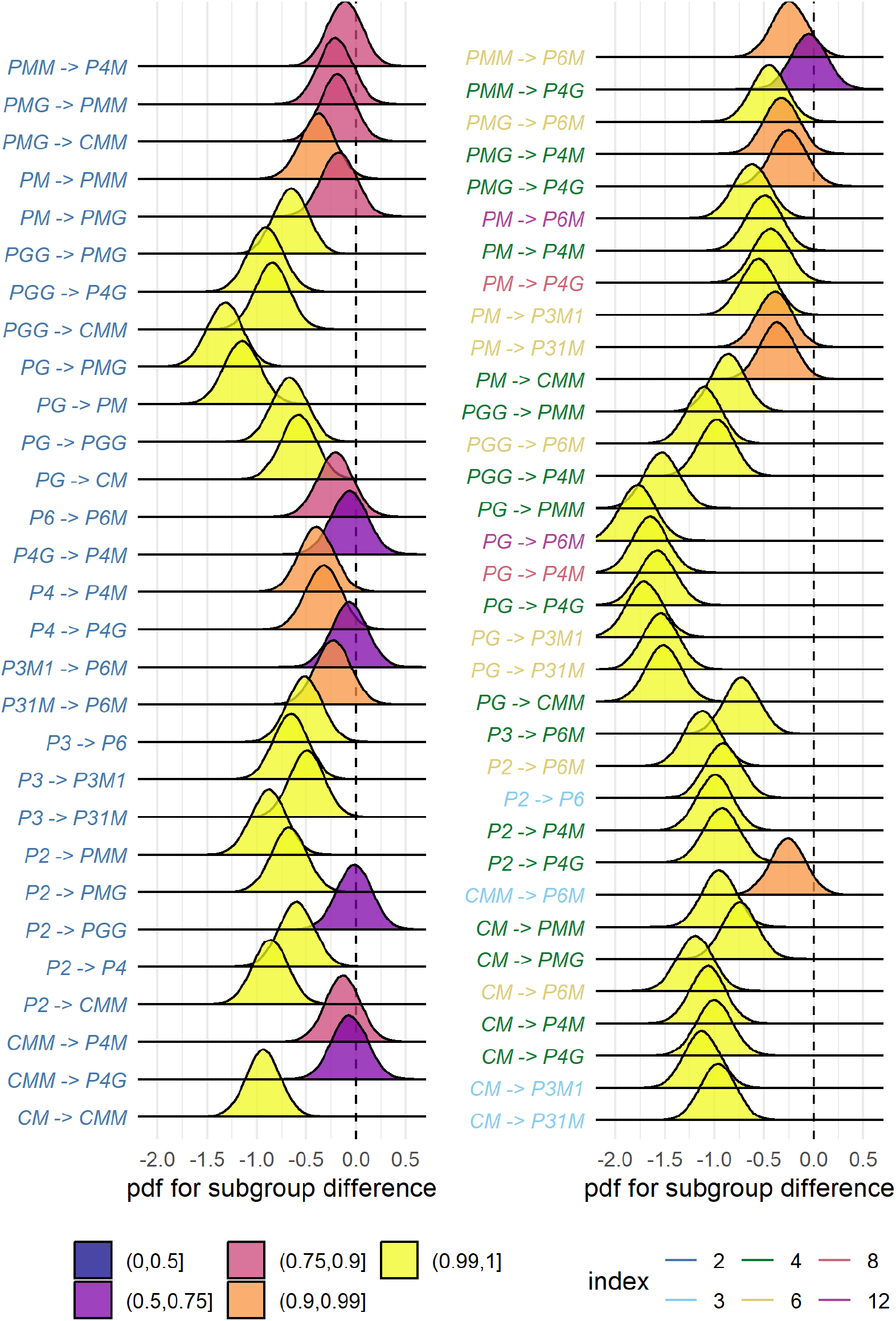
Posterior distributions for the difference in mean symmetry detection threshold dura-tions. Colour coding of the text indicates the index of the subgroup, while the colour of the filled distribution relates to the conditional probability that the difference in means is greater than zero. We can see that 43/63 subgroup relationships have *p*(Δ|*data*) > 0.99.

We also ran a control analysis using (1) odd-harmonic SSVEP data from a six-electrode ROI over parietal cortex (see Supplementary Figure 1.1) and (2) even-harmonic SSVEP data from the same occipital ROI that was used in our primary analysis. By comparing these two control analysis to our primary SSVEP analysis, we can address the specify of our effects in terms of location (occipital cortex vs parietal cortex) and harmonic (odd vs even). For both control analyses (plotted in Supplementary Figures 3.3 and 3.4), the correspondence between data and subgroup relationships was substantially weaker than in the primary analysis. We can quantify the strength of the association between the data and the subgroup relationships, by asking what proportion of subgroup relationships that reach or exceed a range of probability thresholds. This is plotted in Figure 4, for our psychophysical data, our primary SSVEP analysis and our two control SSVEP analyses. It shows that odd-harmonic SSVEP data from the occipital ROI and symmetry detection threshold durations both have a strong association with the subgroup relationships such that a clear majority of the subgroups survive even at the highest threshold we consider (*p*(Δ > O|*data*) > 0.99). The association is far weaker for the two control analyses.

**Figure 4:**
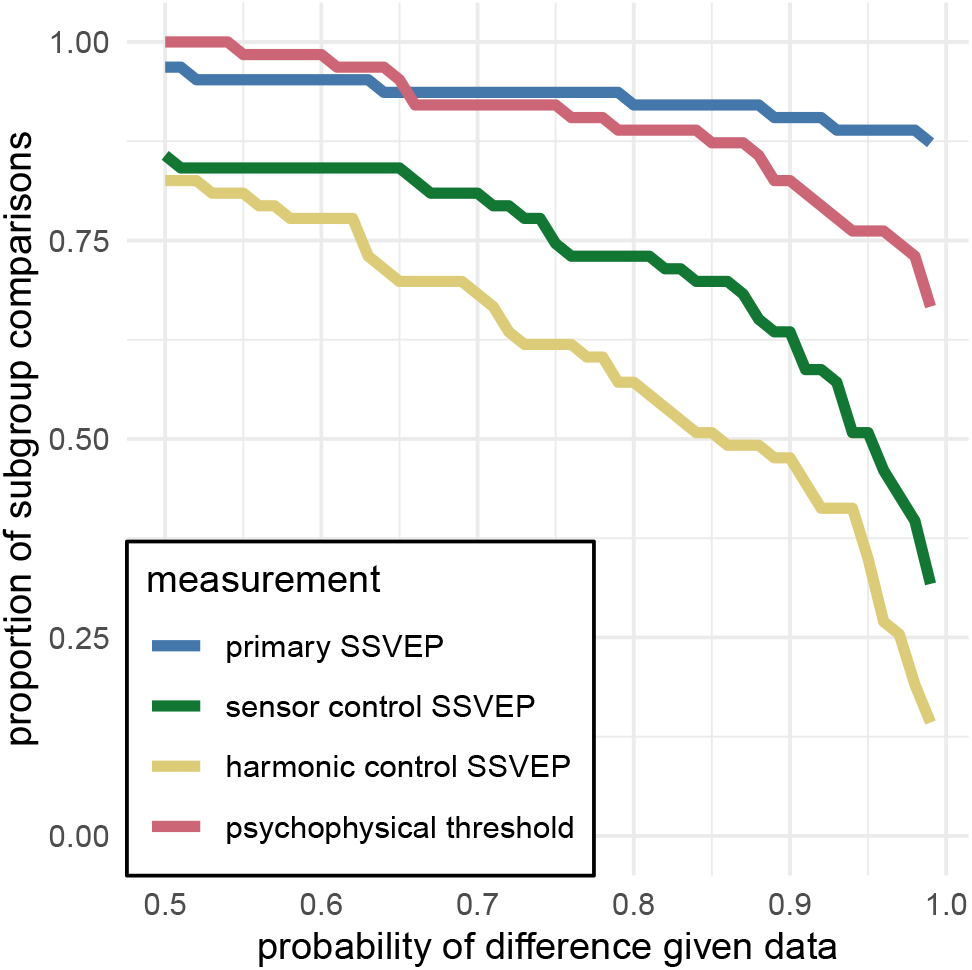
This plot shows the proportion of subgroup relationships that satisfy *p*(Δ > O|*data*) > *x*. We can see that if we take *x* = 0.95 as our threshold, the subgroup relationships are preserved in 56/63 = 89% and 49/64 = 78% of the comparisons for the primary SSVEP and threshold duration datasets, repectively. This compares to the 32/64= 50% and 22/64 = 35% for the SSVEP control datasets.

SSVEP data from four of the wallpaper groups (*P2, P3, P4* and *P6*) was previously published as part of our earlier demonstration of parametric responses to rotation symmetry in wallpaper groups(Kohler et al., 2016). We replicate that result using our Bayesian approach, and find an analogous parametric effect in the psychophysical data (see Supplementary Figure 4.1). We also conducted an analysis testing for an effect of index in our two datasets and found that subgroup relationships with higher indices tended to produce greater pairwise differences between the subgroup and supergroup, for both SSVEP RMS and symmetry detection thresholds (see Supplementary Figure 4.2). The effect of index is relatively weak, but the fact that there is a measurable index effect can nonetheless be taken as preliminary evidence that representations of symmetries in wallpaper groups may be compositional.

Finally, we conducted a correlation analysis comparing SSVEP and psychophysical data and found a reliable correlation (*R*^2^ = 0.44, Bayesian confidence interval [0.28, 0.55]). The correlation reflects an inverse relationship: For subgroup relationships where the supergroup produces a much *larger* SSVEP amplitude than the subgroup, the supergroup also tends to produce a much *smaller* symmetry detection threshold. This is consistent with our hypotheses about how the two measurements relate to symmetry representations in the brain, and suggests that our brain imaging and psychophysical measurements are at least to some extent tapping into the same underlying mechanisms.

## Discussion

Here we show that beyond merely responding to the elementary symmetry operations of reflection (Sasaki et al., 2005) and rotation (Kohler et al., 2016), the visual system represents the hierarchical structure of the 17 wallpaper groups, and thus every composition of the four fundamental symmetries (rotation, reflection, translation, glide reflection) which comprise the set of regular textures. Both SSVEP amplitudes and symmetry detection thresholds preserve the hierarchy of complexity among the wallpaper groups that is captured by the subgroup relationships (Coxeter and Moser, 1972). For the SSVEP, this remarkable consistency was specific to the odd harmonics of the stimulus frequency that are known to capture the symmetry-specific response (Kohler et al., 2016) and to electrodes in a region-of-interest (ROI) over occipital cortex. When the same analysis was done using the odd harmonics from electrodes over parietal cortex (Supplementary Figure 3.3) or even harmonics from electrodes over occipital cortex (Supplementary Figure 3.4), the data was substantially less consistent with the subgroup relationships (yellow and green lines, Figure 4).

The current data provide a description of the visual system’s response to the complete set of symmetries in the two-dimensional plane. Our design precludes us from independently measure the response to *P1*, but because each of the 16 other groups produce non-zero odd harmonic amplitudes (see Figure 1), we can conclude that the relationships between *P1* and all other groups, where *P1* is the subgroup, are also preserved by the visual system. The subgroup relationships are in many cases not obvious perceptually, and most participants had no knowledge of group theory. Thus, the visual system’s ability to preserve the subgroup hierarchy does not depend on explicit knowledge of the relationships. Previous behavioral experiments have shown that although naïve observers can distinguish many of the wallpaper groups (Landwehr, 2009), they tend to sort them into fewer groups than there actually are (4-12 groups) and it is common for exemplars from different wallpaper groups to be sorted in the same group (Clarke et al., 2011). The more controlled two-interval forced choice approach used in the current behavioral experiment allows us to show that more granular representations of wallpaper groups are measurable in behavior.

We observe a reliable correlation between our brain imaging and psychophysical data. This suggests that the two measurements reflect the same underlying symmetry representations in visual cortex. While it should be noted that the correlation is relatively modest (*R*^2^ = 0.44), we note that this may be partly due to the fact that the same individuals did not participate in the two experiments. Future work in which behavioral and brain imaging data are collected from the same participants, will help further establish the connection between the two measurements, and tease apart any additional complexity that may not have been captured by the summary statistics we applied here. It has recently been demonstrated that *W*, a measure of perceptual goodness derived from a holographic model of regularity (van der Helm and Leeuwenberg, 1996), can predict EEG responses (Makin et al., 2016) and perceptual discrimination performance (Nucci and Wagemans, 2007) for patterns that contain symmetry and other types of regularity. The model was formulated based on dot patterns with symmetry axes centered on a single spatial location. It will be important to determine if and how *W* can computed for our random-noise based wallpaper textures where combinations of symmetries tile the plane, and test how well it can explain behavioral and brain responses to wallpapers.

We also find an effect of index for both our brain imaging measurements and our symmetry detection thresholds. This means that the visual system not only represents the hierarchical relationship captured by individual subgroups, but also distinguishes between subgroups depending on how many times the subgroup is repeated in the supergroup, with more repetitions leading to larger pairwise differences. Our measured effect of index is relatively weak. This is perhaps because the index analysis does not take into account the *type* of isometries that differentiate the subgroup and supergroup. The effect of symmetry type can be observed by contrasting the measured SSVEP amplitudes and detection thresholds for groups *PM* and *PG* in Figure 1. The two groups are comparable except *PM* contains reflection and *PG* contains glide reflection, and the former clearly elicits higher amplitudes and lower thresholds. An important goal for future work will be to map out how different symmetry types contribute to the representational hierarchy.

The correspondence between responses in the visual system and group theory that we demonstrate here, may reflect a form of implicit learning that depends on the structure of the natural world. The environment is itself constrained by physical forces underlying pattern formation and these forces are subject to multiple symmetry constraints (Hoyle, 2006). The ordered structure of responses to wallpaper groups could be driven by a central tenet of neural coding, that of efficiency. If coding is to be efficient, neural resources should be distributed to capture the structure of the environment with minimum redundancy considering the visual geometric optics, the capabilities of the subsequent neural coding stages and the behavioral goals of the organism (Attneave, 1954; Barlow, 1961; Laughlin, 1981; Geisler et al., 2009). Early work within the efficient coding framework suggested that natural images had a 1/*f* spectrum and that the corresponding redundancy between pixels in natural images could be coded efficiently with a sparse set of oriented filter responses, such as those present in the early visual pathway (Field, 1987; Olshausen and Field, 1997). Our results suggest that the principle of efficient coding extends to a much higher level of structural redundancy – that of symmetries in visual images.

The 17 wallpaper groups are completely regular, and relatively rare in the visual environment, especially when considering distortions due to perspective and occlusion. Near-regular textures, however, abound in the visual world, and can be approximated as deformed versions of the wallpaper groups (Liu et al., 2004). The correspondence between visual cortex responses and group theory demonstrated here may indicate that the visual system represents visual textures using a similar scheme, with the wallpaper groups serving as anchor points in representational space. This framework resembles norm-based encoding strategies that have been proposed for other stimulus classes, most notably faces (Leopold et al., 2006), and leads to the prediction that adaptation to wallpaper patterns should distort perception of near-regular textures, similar to the aftereffects found for faces (Webster and MacLin, 1999). Field biologists have demonstrated that animals respond more strongly to exaggerated versions of a learned stimulus, referred to as “supernormal” stimuli (Tinbergen, 1953). In the norm-based encoding framework, wallpaper groups can be considered *supertextures*, exaggerated examples of the near-regular textures that surround us. Artists may consciously or unconsciously create supernormal stimuli, to capture the essence of the subject and evoke strong responses in the audience (Ramachandran and Hirstein, 1999). Wallpaper groups are visually compelling, and symmetries have been widely used in human artistic expression going back to the Neolithic age (Jablan, 2014). If wallpapers are in fact supertextures, this prevalence may be a direct result of the strategy the human visual system has adopted for texture encoding.

### Participants

Twenty-five participants (11 females, mean age 28.7±i3.3) took part in the EEG experiment. Their informed consent was obtained before the experiment under a protocol that was approved by the Institutional Review Board of Stanford University. 11 participants (8 females, mean age 2o.73±i.2i) took part in the psychophysics experiment. All participants had normal or corrected-to-normal vision. Their informed consent was obtained before the experiment under a protocol that was approved by the University of Essex’s Ethics Committee.

### Stimulus Generation

Exemplars from the different wallpaper groups were generated using a modified version of the methodology developed by Clarke and colleagues(Clarke et al., 2011) that we have described in detail elsewhere(Kohler et al., 2016). Briefly, exemplar patterns for each group were generated from random-noise textures, which were then repeated and transformed to cover the plane, according to the symmetry axes and geometric lattice specific to each group. The use of noise textures as the starting point for stimulus generation allowed the creation of an almost infinite number of distinct exemplars of each wallpaper group. To make individual exemplars as similar as possible we replaced the power spectrum of each exemplar with the median across exemplars within a group. We then generated control exemplars that had the same power spectrum as the exemplar images by randomizing the phase of each exemplar image. The phase scrambling eliminates rotation, reflection and glide-reflection symmetries within each exemplar, but the phase-scrambled images inherent the spectral periodicity arising from the periodic tiling. This means that all control exemplars, regardless of which wallpaper group they are derived from, are transformed into another symmetry group, namely *P1. P1* is the simplest of the wallpaper groups and contains only translations of a region whose shape derives from the lattice. Because the different wallpaper groups have different lattices, *P1* controls matched to different groups have different power spectra. Our experimental design takes these differences into account by comparing the neural responses evoked by each wallpaper group to responses evoked by the matched control exemplars.

### Stimulus Presentation

Stimulus Presentation. For the EEG experiment, the stimuli were shown on a 24.5” Sony Trimaster EL PVM-2541 organic light emitting diode (OLED) display at a screen resolution of 1920×1080 pixels, 8-bit color depth and a refresh rate of 60 Hz, viewed at a distance of 70 cm. The mean luminance was 69.93 cd/m2 and contrast was 95%. The diameter of the circular aperture in which the wallpaper pattern appeared was 13.8° of visual angle presented against a mean luminance gray background. Stimulus presentation was controlled using in-house software. For the psychophysics experiment, the stimuli were shown on a 48 × 27cm VIEWPixx/3D LCD Display monitor, model VPX-VPX-2005C, resolution 1920 × 1080 pixels, with a viewing distance of approximately 40cm and linear gamma. Stimulus presentation was controlled using MatLab and Psycht00lb0x-3 (Kleiner et al., 2007; Brainard, 1997). The diameter of the circular aperture for the stimuli was 21.5°.

### EEG Procedure

Visual Evoked Potentials were measured using a steady-state design, in which *P1* control images alternated with exemplar images from each of the 16 other wallpaper groups. Exemplar images were always preceded by their matched *P1* control image. A single 0.83 Hz stimulus cycle consisted of a control *P1* image followed by an exemplar image, each shown for 600 ms. A trial consisted of 10 such cycles (12 sec) over which 10 different exemplar images and matched controls from the same rotation group were presented. For each group type, the individual exemplar images were always shown in the same order within the trials. Participants initiated each trial with a button-press, which allowed them to take breaks between trials. Trials from a single wallpaper group were presented in blocks of four repetitions, which were themselves repeated twice per session, and shown in random order within each session. To control fixation, the participants were instructed to fixate a small white cross in the center of display. To control vigilance, a contrast dimming task was employed. Two times per trial, an image pair was shown at reduced contrast, and the participants were instructed to press a button on a response pad. We adjusted the contrast reduction such that average accuracy for each participant was kept at 85% correct, in order to keep the difficulty of the vigilance at a constant level.

### Psychophysics Procedure

The experiment consisted of 16 blocks, one for each of the wallpaper groups (excluding *P1*). We used a two-interval forced choice approach. In each trial, participants were presented with two stimuli (one of which was the wallpaper group for the current block of trials, the other being *P1*), one after the other (inter-stimulus interval of 700ms). After each stimulus had been presented, it was masked with white noise for 300ms. After both stimuli had been presented, participants made a response on the keyboard to indicate whether they thought the first or second image contained more symmetry. Each block started with 10 practice trials, (stimulus display duration of 500ms) to allow participants to familiarise themselves with the current block’s wallpaper pattern. If they achieved an accuracy of 9/10 in these trials they progressed to the rest of the block, otherwise they carried out another set of 10 practise trials. This process was repeated until the required accuracy of 9/10 was obtained. The rest of the block consisted of four interleaved staircases (using the QUEST algorithm (Watson and Pelli, 1983), full details given in the SI) of 30 trials each. On average, a block of trials took around 10 minutes to complete.

### EEG Acquisition and Preprocessing

Steady-State Visual Evoked Potentials (SSVEPs) were collected with 128-sensor HydroCell Sensor Nets (Electrical Geodesics, Eugene, OR) and were band-pass filtered from 0.3 to 50 Hz. Raw data were evaluated off line according to a sample-by-sample thresholding procedure to remove noisy sensors that were replaced by the average of the six nearest spatial neighbors. On average, less than 5% of the electrodes were substituted; these electrodes were mainly located near the forehead or the ears. The substitutions can be expected to have a negligible impact on our results, as the majority of our signal can be expected to come from electrodes over occipital, temporal and parietal cortices. After this operation, the waveforms were re-referenced to the common average of all the sensors. The data from each 12s trial were segmented into five 2.4 s long epochs (i.e., each of these epochs was exactly 2 cycles of image modulation). Epochs for which a large percentage of data samples exceeding a noise threshold (depending on the participant and ranging between 25 and 50 μV) were excluded from the analysis on a sensor-by-sensor basis. This was typically the case for epochs containing artifacts, such as blinks or eye movements. Steady-state stimulation will drive cortical responses at specific frequencies directly tied to the stimulus frequency. It is thus appropriate to quantify these responses in terms of both phase and amplitude. Therefore, a Fourier analysis was applied on every remaining epoch using a discrete Fourier transform with a rectangular window. The use of two-cycle long epochs (i.e., 2.4 s) was motivated by the need to have a relatively high resolution in the frequency domain, δf = 0.42 Hz. For each frequency bin, the complex-valued Fourier coefficients were then averaged across all epochs within each trial. Each participant did two sessions of 8 trials per condition, which resulted in a total of 16 trials per condition.

### SSVEP Analysis

Response waveforms were generated for each group by selective filtering in the frequency domain. For each participant, the average Fourier coefficients from the two sessions were averaged over trials and sessions. The SSVEP paradigm we used allowed us to separate symmetry-related responses from non-specific contrast transient responses. Previous work has demonstrated that symmetry-related responses are predominantly found in the odd harmonics of the stimulus frequency, whereas the even harmonics consist mainly of responses unrelated to symmetry, that arise from the contrast change associated with the appearance of the second image (Norcia et al., 2002; Kohler et al., 2016). This functional distinction of the harmonics allowed us to generate a single-cycle waveform containing the response specific to symmetry, by filtering out the even harmonics in the spectral domain, and then back-transforming the remaining signal, consisting only of odd harmonics, into the time-domain. For our main analysis, we averaged the odd harmonic single-cycle waveforms within a six-electrode region of interest (ROI) over occipital cortex (electrodes 70, 74, 75, 81, 82, 83). These waveforms, averaged over participants, are shown in Figure 1. The same analysis was done for the even harmonics and for the odd harmonics within a six electrode ROI over parietal cortex (electrodes 53, 54, 61, 78, 79, 86; see Supplementary Figure 1.1). The root-mean square values of these waveforms, for each individual participant, were used to determine whether each of the wallpaper subgroup relationships were preserved in the brain data.

### Defining the list of subgroup relationships

In order to get the complete list of subgroup relationships, we digitized Table 4 from Coxeter (Coxeter and Moser, 1972) (shown in Supplementary Table 1.2). After removing identity relationships (i.e. each group is a subgroup of itself) and the three pairs of wallpapers groups that are subgroups of each other (e.g. *PM* is a subgroup of *CM*, and *CM* is a subgroup of *PM)* we were left with a total of 63 unambiguous subgroups that were included in our analysis.

### Bayesian Analysis of SSVEP and Psychophysical data

Bayesian analysis was carried out using R (v3.6.1) (R Core Team, 2019) with the `brms` package (v2.9.0) (Bürkner, 2017) and rStan (v2.19.2 (Stan Development Team, 2019)). The data from each experiment were modelled using a Bayesian generalised mixed effect model with wallpaper group being treated as a 16-level factor, and random effects for participant. The SSVEP data and symmetry detection threshold durations were modelled using log-normal distributions with weakly informative, 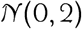, priors. After fitting the model to the data, samples were drawn from the posterior distribution of the two datasets, for each wallpaper group. These samples were then recombined to calculate the distribution of differences for each of the 63 pairs of subgroup and supergroup. These distributions were then summarised by computing the conditional probability of obtaining a positive (negative) difference, *p*(Δ|data). For further technical details, please see the Supplementary Materials at https://osf.io/f3ex8/ where the full R code, model specification, prior and posterior predictive checks, and model diagnostics, can be found.

