## Supplementary material for "The human visual system preserves the hierarchy of 2-dimensional pattern regularity": analysis.html

I have set `echo=FALSE` so that most of the code chunks will not display. Please refer to the .Rmd file for source code.

### 1 Importing data

First we import our data and make some summary plots.

#### 1.1 SSVEP Data

Importing the primary SSVEP data set. This is odd-harmonic filtered data from region-of-interest consisting of six electrodes over occipital cortex.

Figure 1.1: SSVEP RMS data from two of the wallpaper groups, P2 and P4M. Odd harmonics are shown in A and B, while even harmonic data are shown in C and D, and occipital and parietal regions of interest are shown in dark and light gray, respectively. The two groups elicit very different response amplitudes for odd harmonics over occipital cortex, but for even harmonics those differences are much less pronounced.

We now import the data. We have three variables: wallpaper group (wg), subject, and root-mean-squared amplitude (rms).

```
## Rows: 400
## Columns: 3
## $ wg      <chr> "P2", "PM", "PG", "CM", "PMM", "PMG", "PGG", "CMM", "P4", "...
## $ subject <chr> "s01", "s01", "s01", "s01", "s01", "s01", "s01", "s01", "s0...
## $ rms     <dbl> 0.4013, 0.6555, 0.5547, 0.7635, 0.9185, 0.7285, 0.4320, 0.6...
```

If we plot the distribution of rms, we can clearly see that it is skewed. Furthermore, as negative rms amplitudes are impossible we will use a `lognormal` distribution to model these data.

Figure 1.2: Data are the root-mean-squaured (rms) amplitudes over the odd-harmonic filtered waveforms.

#### 1.2 Threshold Data

Here we import that data and select the columns that we’re interested in. Threshold gives the required display duration (in seconds) for the two stimuli to allow for accurate discrimination.

```
## Rows: 186
## Columns: 3
## $ subject   <chr> "person10", "person10", "person10", "person10", "person10...
## $ wg        <chr> "CM", "CMM", "P2", "P3", "P31M", "P3M1", "P4", "P4G", "P4...
## $ threshold <dbl> 0.74125, 0.20216, 0.47697, 0.35012, 0.24529, 0.19022, 0.2...
```

As above, a summary of the data. And again, we have a skewed distribution, (with negative display durations being impossible), so we will use also use a `lognormal` distribution to model the behaviour data.

Figure 1.3: Histogram of the display duration thresholds.

#### 1.3 Control Data

In addition to the primary SSVEP data set, we are also importing two control EEG data sets which are (a) even harmonic data from the same occipital electrodes, and (b) odd harmonic data from six parietal electrodes (see Figure 1.1 and the main paper).

#### 1.4 Symmetry Information

Information on the symmetries and subgroups contained within each wallpaper group was obtained from Coxeter & Moses (1980, Generators and Relations
for Discrete Groups) and is summarised in the files `symmeries_in_group.csv`, `subgroup_relations.csv`, and `subgroup_normal.csv`.

##### 1.4.1 Types of Symmetry

The following table lists the non-translational isometries in the 17 wallpaper groups. Isometries are listed up to conjugacy, meaning for instance that two identical rotations about different points are not counted separately, and nor are two reflections about lines which differ by a translation in the wallpaper group.

Table 1.1: Wallpaper group summary

| group | rotation | reflection | glide |
| --- | --- | --- | --- |
| P2 | 2 | 0 | 0 |
| PM | 0 | 1 | 0 |
| PG | 0 | 0 | 1 |
| CM | 0 | 1 | 1 |
| PMM | 2 | 2 | 0 |
| PMG | 2 | 1 | 1 |
| PGG | 2 | 0 | 2 |
| CMM | 2 | 2 | 2 |
| P4 | 4 | 0 | 0 |
| P4M | 4 | 4 | 2 |
| P4G | 4 | 2 | 4 |
| P3 | 3 | 0 | 0 |
| P3M1 | 3 | 3 | 3 |
| P31M | 3 | 3 | 3 |
| P6 | 6 | 0 | 0 |
| P6M | 6 | 6 | 6 |

##### 1.4.2 Subgroup Relationships

Import subgroup information and display a table of the relationships that we will be investigating. Relationships taken from Coxeter & Moser (1972).

Table 1.2: Summary of subgroup relationships. The numbers indicate the index of the subgroup, while italics indicate normal subgroups. Relationships written in yellow text are not included in our analysis.

| subgroup | P2 | PG | PM | CM | PGG | PMG | PMM | CMM | P4 | P4G | P4M | P3 | P3M1 | P31M | P6 | P6M |
| --- | --- | --- | --- | --- | --- | --- | --- | --- | --- | --- | --- | --- | --- | --- | --- | --- |
| P2 | 2 | - | - | - | - | - | - | - | - | - | - | - | - | - | - | - |
| PG | - | 2 | - | - | - | - | - | - | - | - | - | - | - | - | - | - |
| PM | - | 2 | 2 | 2 | - | - | - | - | - | - | - | - | - | - | - | - |
| CM | - | 2 | 2 | 3 | - | - | - | - | - | - | - | - | - | - | - | - |
| PGG | 2 | 2 | - | - | 3 | - | - | - | - | - | - | - | - | - | - | - |
| PMG | 2 | 2 | 2 | 4 | 2 | 3 | - | - | - | - | - | - | - | - | - | - |
| PMM | 2 | 4 | 2 | 4 | 4 | 2 | 2 | 2 | - | - | - | - | - | - | - | - |
| CMM | 2 | 4 | 4 | 2 | 2 | 2 | 2 | 4 | - | - | - | - | - | - | - | - |
| P4 | 2 | - | - | - | - | - | - | - | 2 | - | - | - | - | - | - | - |
| P4G | 4 | 4 | 8 | 4 | 2 | 4 | 4 | 2 | 2 | 9 | - | - | - | - | - | - |
| P4M | 4 | 8 | 4 | 4 | 4 | 4 | 2 | 2 | 2 | 2 | 2 | - | - | - | - | - |
| P3 | - | - | - | - | - | - | - | - | - | - | - | 3 | - | - | - | - |
| P3M1 | - | 6 | 6 | 3 | - | - | - | - | - | - | - | 2 | 4 | 3 | - | - |
| P31M | - | 6 | 6 | 3 | - | - | - | - | - | - | - | 2 | 3 | 4 | - | - |
| P6 | 3 | - | - | - | - | - | - | - | - | - | - | 2 | - | - | 4 | - |
| P6M | 6 | 12 | 12 | 6 | 6 | 6 | 6 | 3 | - | - | - | 4 | 2 | 2 | 2 | 3 |

We will remove identity relationships (i.e., a group is a subgroup of itself) and the three pairs of wallpapers groups that are subgroups of each other (e.g., PM is a subgroup of CM, and CM is a subgroup of PM). This leaves us with a total of 63 subgroups to include in our analysis.

### 2 Bayesian Analysis

Here are the details of the Bayesian multi-level modelling. Our general approach is:

- Use weakly-informative priors.
- Fit a multi-level model using the lognormal distribution.
- Independent variable (wallpaper group) and one dependent variable (rms or threshold).
- Maximal random effect structure.
- Extract mcmc samples from the posterior distributions.
- Use these samples to estimate the difference between sub- and supergroups.

#### 2.1 Define Priors

In this section we will specify some priors. We then then use a prior-predictive check to assess whether the prior is reasonable or not (i.e., on the same order of magnitude as our measurements).

##### 2.1.1 Fixed Effects

Our independent variable is a categorical factor with 16 levels. We will drop the intercept from our model and instead fit a coefficient for each factor level (\(y \sim x - 0\)). As our dependent variable will be log-transformed, we can use the priors below:

```
prior <- c(
  set_prior("normal(0,2)", class = "b"),    
  set_prior("cauchy(0,2)", class = "sigma"))
```

##### 2.1.2 Group-level Effects

We will keep the default weakly informative priors for the group-level (‘random’) effects. From the brms documentation:

> […] restricted to be non-negative and, by default, have a half student-t prior with 3 degrees of freedom and a scale parameter that depends on the standard deviation of the response after applying the link function. Minimally, the scale parameter is 10. This prior is used (a) to be only very weakly informative in order to influence results as few as possible, while (b) providing at least some regularization to considerably improve convergence and sampling efficiency.

##### 2.1.3 Prior Predictive Check

Now we can specify our Bayesian multi-level model and priors. Note that as we are using `sample_prior = 'only'`, the model will not learn anything from our data.

```
m_prior <- brm(data = d_eeg, 
  rms ~ wg-1 + (1|subject),
  family = "lognormal", 
  prior = prior, 
  iter = n_iter ,
  sample_prior = 'only')
```

We can use this model to generate data.

```
## Rows: 320,000
## Columns: 2
## $ key   <chr> "P2", "P2", "P2", "P2", "P2", "P2", "P2", "P2", "P2", "P2", "...
## $ value <dbl> 0.52969256, 0.61739515, 0.08499849, 11.72765850, 0.12273652, ...
```

```
## Warning: 'stat_pointintervalh' is deprecated.
## Use 'stat_pointinterval' instead.
## See help("Deprecated") and help("tidybayes-deprecated").

## Warning: 'stat_pointintervalh' is deprecated.
## Use 'stat_pointinterval' instead.
## See help("Deprecated") and help("tidybayes-deprecated").
```

Figure 2.1: The density plot shows the distribution of the empirical data, while the blue line shows the 66% and 95% prediction intervals.

We can see that i) our priors are relatively weak as the predictions span several orders of magnitude, and ii) our empirical data falls within this range.

#### 2.2 Compute Posterior

##### 2.2.1 Fit Model to SSVEP Data

We will now fit the model to the data.

```
##  Family: lognormal 
##   Links: mu = identity; sigma = identity 
## Formula: rms ~ wg - 1 + (1 | subject) 
##    Data: d_eeg (Number of observations: 400) 
## Samples: 4 chains, each with iter = 10000; warmup = 5000; thin = 1;
##          total post-warmup samples = 20000
## 
## Group-Level Effects: 
## ~subject (Number of levels: 25) 
##               Estimate Est.Error l-95% CI u-95% CI Rhat Bulk_ESS Tail_ESS
## sd(Intercept)     0.38      0.06     0.28     0.52 1.00     1403     2810
## 
## Population-Level Effects: 
##        Estimate Est.Error l-95% CI u-95% CI Rhat Bulk_ESS Tail_ESS
## wgCM      -0.49      0.09    -0.65    -0.32 1.00      764     1288
## wgCMM     -0.13      0.09    -0.29     0.05 1.01      753     1197
## wgP2      -0.95      0.09    -1.12    -0.77 1.01      772     1190
## wgP3      -0.82      0.09    -0.98    -0.65 1.00      771     1200
## wgP31M    -0.44      0.09    -0.60    -0.26 1.00      757     1255
## wgP3M1    -0.14      0.09    -0.31     0.03 1.01      762     1206
## wgP4      -0.49      0.09    -0.65    -0.31 1.00      766     1288
## wgP4G     -0.26      0.09    -0.42    -0.08 1.00      764     1186
## wgP4M      0.15      0.09    -0.02     0.32 1.01      784     1292
## wgP6      -0.48      0.09    -0.65    -0.31 1.01      762     1205
## wgP6M     -0.05      0.09    -0.21     0.13 1.00      787     1307
## wgPG      -0.93      0.09    -1.10    -0.76 1.01      768     1214
## wgPGG     -0.73      0.09    -0.89    -0.55 1.00      761     1222
## wgPM      -0.60      0.09    -0.76    -0.42 1.00      763     1224
## wgPMG     -0.26      0.09    -0.43    -0.09 1.01      736     1249
## wgPMM     -0.07      0.09    -0.23     0.10 1.00      783     1255
## 
## Family Specific Parameters: 
##       Estimate Est.Error l-95% CI u-95% CI Rhat Bulk_ESS Tail_ESS
## sigma     0.23      0.01     0.21     0.25 1.00     9343    11753
## 
## Samples were drawn using sampling(NUTS). For each parameter, Bulk_ESS
## and Tail_ESS are effective sample size measures, and Rhat is the potential
## scale reduction factor on split chains (at convergence, Rhat = 1).
```

We will now look at the model’s predicts for the average participant (i.e, ignoring the random intercepts).

```
## Warning: 'stat_pointintervalh' is deprecated.
## Use 'stat_pointinterval' instead.
## See help("Deprecated") and help("tidybayes-deprecated").

## Warning: 'stat_pointintervalh' is deprecated.
## Use 'stat_pointinterval' instead.
## See help("Deprecated") and help("tidybayes-deprecated").
```

Figure 2.2: The density plot shows the distribution of the empirical data, while the red line shows the 66% and 95% prediction intervals.

##### 2.2.2 Fit Model to Psychophysical Data

We will now fit the model to the data.

```
##  Family: lognormal 
##   Links: mu = identity; sigma = identity 
## Formula: threshold ~ wg - 1 + (1 | subject) 
##    Data: d_dispthresh (Number of observations: 186) 
## Samples: 4 chains, each with iter = 10000; warmup = 5000; thin = 1;
##          total post-warmup samples = 20000
## 
## Group-Level Effects: 
## ~subject (Number of levels: 12) 
##               Estimate Est.Error l-95% CI u-95% CI Rhat Bulk_ESS Tail_ESS
## sd(Intercept)     0.39      0.11     0.23     0.65 1.00     3459     6092
## 
## Population-Level Effects: 
##        Estimate Est.Error l-95% CI u-95% CI Rhat Bulk_ESS Tail_ESS
## wgCM      -0.22      0.17    -0.54     0.11 1.00     3142     5634
## wgCMM     -1.16      0.16    -1.47    -0.83 1.00     3196     5370
## wgP2      -0.29      0.17    -0.62     0.05 1.00     3222     5645
## wgP3      -0.69      0.16    -1.01    -0.37 1.00     3069     5387
## wgP31M    -1.19      0.17    -1.50    -0.85 1.00     3097     5384
## wgP3M1    -1.35      0.17    -1.67    -1.01 1.00     3264     5740
## wgP4      -0.89      0.17    -1.21    -0.56 1.00     3095     5213
## wgP4G     -1.22      0.17    -1.55    -0.88 1.00     3304     5759
## wgP4M     -1.29      0.17    -1.61    -0.96 1.00     3157     5012
## wgP6      -1.21      0.17    -1.53    -0.87 1.00     3196     5815
## wgP6M     -1.42      0.17    -1.74    -1.08 1.00     3106     5526
## wgPG       0.36      0.17     0.04     0.70 1.00     3246     5553
## wgPGG     -0.31      0.17    -0.63     0.02 1.00     3168     5469
## wgPM      -0.79      0.17    -1.11    -0.45 1.00     3132     5513
## wgPMG     -0.97      0.17    -1.29    -0.64 1.00     3156     5794
## wgPMM     -1.18      0.17    -1.49    -0.85 1.00     2956     5477
## 
## Family Specific Parameters: 
##       Estimate Est.Error l-95% CI u-95% CI Rhat Bulk_ESS Tail_ESS
## sigma     0.42      0.02     0.37     0.46 1.00    14199    13704
## 
## Samples were drawn using sampling(NUTS). For each parameter, Bulk_ESS
## and Tail_ESS are effective sample size measures, and Rhat is the potential
## scale reduction factor on split chains (at convergence, Rhat = 1).
```

##### 2.2.3 EEG Control Data

We will also fit models to the control data. As we can see from Figure 2.4, the group differences are much smaller.

Figure 2.3: The density plot shows the distribution of the empirical data, while the blue line shows the 66% and 95% prediction intervals.

### 3 Subgroup Comparisons

We will now compute the difference between sub- and supergroups.

#### 3.1 Primary SSVEP Data

Finally, we calculate the probability that the SSVEP RMS difference between subgroup and supergroup is larger than zero given the data. This information is then binned so we can colour in the posterior density plots.

Figure 3.1: Distributions of the difference in mean log(rms) between sub- and supergroups. The index of each relationship is indicated by the colour of the y-axis label. The fill of the density plots indicated the probability of the difference being greater than zero.

#### 3.2 Psychophysical Data

We can do the same for the display duration thresholds from our psychophysics experiment. Here we are looking for the opposite effect, namely that display larger are larger for subgroups than for supergroups (see main paper), so we calculate the probability that differences in duration are smaller than zero.

Figure 3.2: Distributions of the difference in mean log display duration threshold between sub- and supergroups. The index of each relationship is indicated by the colour of the y-axis label. The fill of the density plots indicated the probability of the difference being less than zero.

#### 3.3 Control SSVEP Data

We will now do exactly the same with the control data (odd harmonic data from parietal electrodes and even harmonic data from occipital electrodes)

Figure 3.3: Distributions of the difference in mean log(rms) between sub- and supergroups. The index of each relationship is indicated by the colour of the y-axis label. The fill of the density plots indicated the probability of the difference being greater than zero.

Figure 3.4: Distributions of the difference in mean log(rms) between sub- and supergroups. The index of each relationship is indicated by the colour of the y-axis label. The fill of the density plots indicated the probability of the difference being greater than zero.

#### 3.4 Summary

We can summarise the subgroup comparison plots above by plotting ROC curves for each of our four measurements (Figure 3.5).

| eeg | threshold | occ\_even | par\_odd | p |
| --- | --- | --- | --- | --- |
| 56 | 48 | 32 | 22 | 0.95 |

Figure 3.5: This figure shows how many of our 63 comparisons are classed as having a greater-than-zero difference (less-than-zero for the display durations) for difference thresholds. between 0.5 an 1.0.

If we take \(p\)=0.95 as our cut-off, we can see that the subgroup relations are preserved in 56/63 = 89% and 49/63 = 78% of the comparisons for the primary SSVEP and display durations respectively. This compares to the 32/63= 51% and 22/63 = 35% for the control SSVEP conditions.

### 4 Additional Analysis

#### 4.1 Replication of Kohler et al (2016)

We can look at the groups that only contain rotations, and see if we obtain the parametric response as documented in Kohler et al. (2016).

```
##  Family: lognormal 
##   Links: mu = identity; sigma = identity 
## Formula: rms ~ rotation + (1 | subject) 
##    Data: d_eeg_rot (Number of observations: 100) 
## Samples: 4 chains, each with iter = 2000; warmup = 1000; thin = 1;
##          total post-warmup samples = 4000
## 
## Group-Level Effects: 
## ~subject (Number of levels: 25) 
##               Estimate Est.Error l-95% CI u-95% CI Rhat Bulk_ESS Tail_ESS
## sd(Intercept)     0.39      0.07     0.28     0.54 1.00     1052     1803
## 
## Population-Level Effects: 
##           Estimate Est.Error l-95% CI u-95% CI Rhat Bulk_ESS Tail_ESS
## Intercept    -1.15      0.11    -1.36    -0.94 1.00     1348     2198
## rotation      0.12      0.02     0.09     0.16 1.00     6191     2891
## 
## Family Specific Parameters: 
##       Estimate Est.Error l-95% CI u-95% CI Rhat Bulk_ESS Tail_ESS
## sigma     0.27      0.02     0.23     0.32 1.00     3525     3171
## 
## Samples were drawn using sampling(NUTS). For each parameter, Bulk_ESS
## and Tail_ESS are effective sample size measures, and Rhat is the potential
## scale reduction factor on split chains (at convergence, Rhat = 1).
```

We will also investigate if we see the corresponding pattern with the display duration threshold data, with the time taken to detect the symmetry decreasing as we increase the amount of rotational symmetry.

```
##  Family: lognormal 
##   Links: mu = identity; sigma = identity 
## Formula: threshold ~ rotation + (1 | subject) 
##    Data: d_disp_rot (Number of observations: 46) 
## Samples: 4 chains, each with iter = 2000; warmup = 1000; thin = 1;
##          total post-warmup samples = 4000
## 
## Group-Level Effects: 
## ~subject (Number of levels: 12) 
##               Estimate Est.Error l-95% CI u-95% CI Rhat Bulk_ESS Tail_ESS
## sd(Intercept)     0.47      0.17     0.18     0.86 1.00      995     1505
## 
## Population-Level Effects: 
##           Estimate Est.Error l-95% CI u-95% CI Rhat Bulk_ESS Tail_ESS
## Intercept    -0.01      0.25    -0.50     0.48 1.00     2091     2026
## rotation     -0.22      0.05    -0.31    -0.12 1.00     4129     2359
## 
## Family Specific Parameters: 
##       Estimate Est.Error l-95% CI u-95% CI Rhat Bulk_ESS Tail_ESS
## sigma     0.47      0.06     0.37     0.62 1.00     1735     2114
## 
## Samples were drawn using sampling(NUTS). For each parameter, Bulk_ESS
## and Tail_ESS are effective sample size measures, and Rhat is the potential
## scale reduction factor on split chains (at convergence, Rhat = 1).
```

Figure 4.1: Red lines show empirical data, blue lines show the model fit.

#### 4.2 Index

Subgroup relations can be classified by their index. Here we investigate the extent to which index can account for the variation between the subgroup relationships.

First of all, we run for the SSVEP RMS data.

```
##  Family: gaussian 
##   Links: mu = identity; sigma = identity 
## Formula: mean_value ~ index 
##    Data: comp_summary$eeg (Number of observations: 63) 
## Samples: 4 chains, each with iter = 2000; warmup = 1000; thin = 1;
##          total post-warmup samples = 4000
## 
## Population-Level Effects: 
##           Estimate Est.Error l-95% CI u-95% CI Rhat Bulk_ESS Tail_ESS
## Intercept     0.34      0.07     0.20     0.48 1.00     3403     2471
## index         0.03      0.02     0.00     0.07 1.00     3329     2406
## 
## Family Specific Parameters: 
##       Estimate Est.Error l-95% CI u-95% CI Rhat Bulk_ESS Tail_ESS
## sigma     0.29      0.03     0.24     0.35 1.00     3439     2836
## 
## Samples were drawn using sampling(NUTS). For each parameter, Bulk_ESS
## and Tail_ESS are effective sample size measures, and Rhat is the potential
## scale reduction factor on split chains (at convergence, Rhat = 1).
```

And now for the display duration thresholds.

```
##  Family: gaussian 
##   Links: mu = identity; sigma = identity 
## Formula: mean_value ~ index 
##    Data: comp_summary$threshold (Number of observations: 63) 
## Samples: 4 chains, each with iter = 2000; warmup = 1000; thin = 1;
##          total post-warmup samples = 4000
## 
## Population-Level Effects: 
##           Estimate Est.Error l-95% CI u-95% CI Rhat Bulk_ESS Tail_ESS
## Intercept    -0.40      0.11    -0.62    -0.18 1.00     3607     2637
## index        -0.09      0.03    -0.13    -0.03 1.00     3786     2939
## 
## Family Specific Parameters: 
##       Estimate Est.Error l-95% CI u-95% CI Rhat Bulk_ESS Tail_ESS
## sigma     0.44      0.04     0.37     0.53 1.00     3503     2511
## 
## Samples were drawn using sampling(NUTS). For each parameter, Bulk_ESS
## and Tail_ESS are effective sample size measures, and Rhat is the potential
## scale reduction factor on split chains (at convergence, Rhat = 1).
```

We can see that the index of the subgroup relationship has an effect on both the difference in log(rms) and the difference in log(display duration): relationships with a higher index lead to larger differences.

Figure 4.2: The effect of index on log(rms) and log(ms).

#### 4.3 Correlation Between Primary SSVEP data and Psychophysical Thresholds

Finally, we will investigate whether there is a correlation between the our primary SSVEP measure (RMS amplitude of odd harmonics over occipital cortex) and our display duration thresholds. As our two different measures come from different samples of participants, we are unable to do a direct comparison. However, we can use the results of the models discussed in Section 3 and check for a correlation between the predicted values of the two measures.

```
##     Estimate  Est.Error      Q2.5     Q97.5
## R2 0.4375808 0.07245691 0.2787564 0.5552389
```

The confidence interval indicates that we can be confident that \(R^2>0\) (i.e, 95% credible interval is 0.28 - 0.56).

Figure 4.3: Scatter plot showing the correlation between our two measures. Each line is a sample from the posterior of a Bayesian linear regression.

### 5 Package and Session Information

Details of packages, etc, are given below.

```
## R version 4.0.3 (2020-10-10)
## Platform: x86_64-w64-mingw32/x64 (64-bit)
## Running under: Windows 10 x64 (build 19041)
## 
## Matrix products: default
## 
## locale:
## [1] LC_COLLATE=English_United Kingdom.1252 
## [2] LC_CTYPE=English_United Kingdom.1252   
## [3] LC_MONETARY=English_United Kingdom.1252
## [4] LC_NUMERIC=C                           
## [5] LC_TIME=English_United Kingdom.1252    
## 
## attached base packages:
## [1] stats     graphics  grDevices utils     datasets  methods   base     
## 
## other attached packages:
##  [1] magick_2.6.0    patchwork_1.1.1 ggthemes_4.2.4  see_0.6.1      
##  [5] latex2exp_0.4.0 ggridges_0.5.3  tidybayes_2.3.1 brms_2.14.4    
##  [9] Rcpp_1.0.6      forcats_0.5.1   stringr_1.4.0   dplyr_1.0.4    
## [13] purrr_0.3.4     readr_1.4.0     tidyr_1.1.2     tibble_3.0.6   
## [17] ggplot2_3.3.3   tidyverse_1.3.0 bookdown_0.21  
## 
## loaded via a namespace (and not attached):
##   [1] readxl_1.3.1         backports_1.2.1      plyr_1.8.6          
##   [4] igraph_1.2.6         splines_4.0.3        svUnit_1.0.3        
##   [7] crosstalk_1.1.1      rstantools_2.1.1     inline_0.3.17       
##  [10] digest_0.6.27        htmltools_0.5.1.1    rsconnect_0.8.16    
##  [13] fansi_0.4.2          magrittr_2.0.1       modelr_0.1.8        
##  [16] RcppParallel_5.0.2   matrixStats_0.58.0   xts_0.12.1          
##  [19] prettyunits_1.1.1    colorspace_2.0-0     rvest_0.3.6         
##  [22] ggdist_2.4.0         haven_2.3.1          xfun_0.20           
##  [25] callr_3.5.1          crayon_1.4.0         jsonlite_1.7.2      
##  [28] lme4_1.1-26          zoo_1.8-8            glue_1.4.2          
##  [31] kableExtra_1.3.1     gtable_0.3.0         webshot_0.5.2       
##  [34] V8_3.4.0             distributional_0.2.2 pkgbuild_1.2.0      
##  [37] rstan_2.21.2         abind_1.4-5          scales_1.1.1        
##  [40] mvtnorm_1.1-1        DBI_1.1.1            miniUI_0.1.1.1      
##  [43] viridisLite_0.3.0    xtable_1.8-4         ggstance_0.3.5      
##  [46] stats4_4.0.3         StanHeaders_2.21.0-7 DT_0.17             
##  [49] htmlwidgets_1.5.3    httr_1.4.2           threejs_0.3.3       
##  [52] arrayhelpers_1.1-0   ellipsis_0.3.1       pkgconfig_2.0.3     
##  [55] loo_2.4.1            farver_2.0.3         dbplyr_2.1.0        
##  [58] utf8_1.1.4           labeling_0.4.2       tidyselect_1.1.0    
##  [61] rlang_0.4.10         reshape2_1.4.4       later_1.1.0.1       
##  [64] effectsize_0.4.3     munsell_0.5.0        cellranger_1.1.0    
##  [67] tools_4.0.3          cli_2.3.0            generics_0.1.0      
##  [70] broom_0.7.4          evaluate_0.14        fastmap_1.1.0       
##  [73] yaml_2.2.1           processx_3.4.5       knitr_1.31          
##  [76] fs_1.5.0             nlme_3.1-149         mime_0.9            
##  [79] projpred_2.0.2       xml2_1.3.2           compiler_4.0.3      
##  [82] bayesplot_1.8.0      shinythemes_1.2.0    rstudioapi_0.13     
##  [85] gamm4_0.2-6          curl_4.3             reprex_1.0.0        
##  [88] statmod_1.4.35       stringi_1.5.3        highr_0.8           
##  [91] ps_1.5.0             parameters_0.11.0    Brobdingnag_1.2-6   
##  [94] lattice_0.20-41      Matrix_1.2-18        nloptr_1.2.2.2      
##  [97] markdown_1.1         shinyjs_2.0.0        vctrs_0.3.6         
## [100] pillar_1.4.7         lifecycle_0.2.0      bridgesampling_1.0-0
## [103] cowplot_1.1.1        insight_0.12.0       httpuv_1.5.5        
## [106] R6_2.5.0             promises_1.1.1       gridExtra_2.3       
## [109] codetools_0.2-16     boot_1.3-25          colourpicker_1.1.0  
## [112] MASS_7.3-53          gtools_3.8.2         assertthat_0.2.1    
## [115] withr_2.4.1          shinystan_2.5.0      mgcv_1.8-33         
## [118] bayestestR_0.8.2     parallel_4.0.3       hms_1.0.0           
## [121] grid_4.0.3           coda_0.19-4          minqa_1.2.4         
## [124] rmarkdown_2.6        shiny_1.6.0          lubridate_1.7.9.2   
## [127] base64enc_0.1-3      dygraphs_1.1.1.6
```
